# annonex2embl: automatic preparation of annotated DNA sequences for bulk submissions to ENA

**DOI:** 10.1101/820480

**Authors:** Michael Gruenstaeudl

## Abstract

**Motivation:** The submission of annotated sequence data to public sequence databases constitutes a central pillar in biological research. The surge of novel DNA sequences awaiting database submission due to the application of next-generation sequencing has increased the need for software tools that facilitate bulk submissions. This need has yet to be met with a concurrent development of tools to automate the preparatory work preceding such submissions.

**Results:** I introduce `annonex2embl`, a Python package that automates the preparation of complete sequence flatfiles for large-scale sequence submissions to the European Nucleotide Archive. The tool enables the conversion of DNA sequence alignments that are co-supplied with sequence annotations and metadata to submission-ready flatfiles. Among other features, the software automatically accounts for length differences among the input sequences while maintaining correct annotations, automatically interlaces metadata to each record, and displays a design suitable for easy integration into bioinformatic workflows. As proof of its utility, `annonex2embl` is employed in preparing a dataset of more than 1,500 fungal DNA sequences for database submission.

## INTRODUCTION

The submission of nucleotide sequence data to public sequence databases is a central pillar in the academic effort to share scientific data and make biological research more reproducible (Tenopir et al. 2011; Drew et al. 2013; Blaxter et al. 2016). In fact, submitting newly generated sequences to public repositories has become a mandatory practice for many scientific journals in the biological sciences (Fairbairn 2011; Vines et al. 2013; Roche et al. 2015). Several sequence databases accept nucleotide sequences for public storage, including the DNA Data Bank of Japan (DDBJ, Kodama et al. 2018), the European Nucleotide Archive (ENA, Harrison et al. 2019), and the nucleotide section of the National Center for Biotechnology Information (NCBI GenBank, Benson et al. 2018). These three archives operate under the umbrella of the International Nucleotide Sequence Database Collaboration (INSDC, Karsch-Mizrachi et al. 2018), which coordinates the standardized storage and public access of submitted sequences and has defined common data standards for the sequences and associated metadata (Cochrane et al. 2015).

The almost ubiquitous implementation of next-generation sequencing in biological research has led to a surge of novel DNA sequences from various organisms (Kress et al. 2015; Levy and Myers 2016) and made the generation and analysis of large-scale biological datasets commonplace (Hampton et al. 2013; Farley et al. 2018). Many contemporary molecular phylogenetic and population genetic studies now generate hundreds, if not thousands, of novel nucleotide sequences per investigation (e.g., Leebens-Mack et al. 2019; Li et al. 2019; Varga et al. 2019; Zhao et al. 2019). Accordingly, a considerable fraction of submissions to public databases represents sequence data from large-scale investigations (Kans and Ouellette 2001; Gruenstaeudl and Hartmaring 2019). Data accumulation statistics for nucleotide sequences at ENA corroborate this trend (Silvester et al. 2018; Cook et al. 2019). Most sequence databases have, consequently, adopted policies and methods to enable the submission of large-scale sequence datasets. GenBank, for example, has implemented a submission workflow to facilitate the sequence upload and submission for targeted locus studies containing 2,500 or more ribosomal RNA sequences (Sayers et al. 2019). It also expanded the functionality of its command-line submission tool (tbl2asn, Benson et al. 2006) to accept sequence data in different file formats (Sayers et al. 2019). Similarly, ENA has implemented measures to automate the sequence upload process through a new command-line submission tool (Webin-CLI, Harrison et al. 2019) and, thus, facilitate the streamlined submission of large numbers of annotated DNA sequences via its Webin submission portal (Silvester et al. 2018). However, the development of software tools necessary to format and prepare DNA sequences for large-scale submissions has not kept pace with the methodological advances to upload them. While command-line driven data transfer procedures can facilitate the upload of an almost infinite number of sequences to public sequence databases, the compilation of the necessary input files is a separate step and also requires automation.

Automating the preparation of novel sequence data for submission to public sequence databases is an important objective and particularly relevant in investigations that generate large numbers of similarly structured nucleotide sequences. For example, molecular phylogenetic and population genetic studies analyze differences in nucleotide sequence data among closely-related taxa (Yang and Rannala 2012; Casillas and Barbadilla 2017) and typically generate DNA sequence alignments (Morrison 2006). Such alignments represent ideal starting points for the automated preparation of submission input files. At that stage, the positional homology among the aligned sequences allows the bulk assignment of functional annotations across all sequences of a dataset, regardless of sequence number or length. Moreover, all sequences of a molecular phylogenetic or population genetic dataset must pass through the process of multiple sequence alignment (MSA) to establish positional homology (Morrison et al. 2015), rendering this stage optimal for the interlacing of sequence metadata. Preparing sequence submissions at the stage of MSA, thus, has important advantages compared to other submission preparation strategies. However, few, if any, software tools can facilitate the automated preparation of aligned and annotated DNA sequences for submission to public sequence databases. To the best of my knowledge, no stand-alone software tool currently exists that can automatically convert a large set of annotated DNA sequences and associated metadata into a file format compatible with the upload requirements of ENA. Bulk sequence submissions to ENA primarily operate with data files formatted according to the EMBL flatfile standard (Silvester et al. 2018); EMBL-formatted flatfiles are compact, yet human-readable, and contain the nucleotide sequences of an organism, functional annotations, and associated metadata (Stoesser et al. 2002). Similarly, no stand-alone file conversion tool currently exists that can generate EMBL formatted flatfiles and simultaneously be integrated into automated bioinformatic pipelines; the software tools available are primarily driven by graphical user interfaces (e.g., Artemis, Rutherford et al. 2000; DnaSP, Rozas et al. 2017) and, thus, mandate researchers to edit sequences by hand. Evidently, a software tool is needed that streamlines and automates the preparatory steps for the submission of large datasets of aligned and annotated DNA sequences to ENA. Specifically, the scientific community would benefit from an open-source, command-line driven software tool that automates the interlacing and conversion of MSAs, sequence annotations, and sequence metadata into submission-ready EMBL flatfiles.

In the present investigation, I introduce a software tool that automates the generation of submission-ready EMBL flatfiles from annotated DNA sequence alignments and associated metadata. The tool, titled `annonex2embl`, parses, splits and re-formats DNA sequences and their associated annotations from an MSA, supplements each sequence with the correct metadata of a co-supplied spreadsheet, and saves each sequence in the EMBL flatfile format. The software contains a series of features for the interlacing of nucleotide sequences, sequence annotations, and sequence metadata, and streamlines the process of preparing DNA sequences from molecular phylogenetic or population genetic investigations for submission to ENA. With the application of `annonex2embl`, a user can automatically generate a multi-record flatfile suitable for the bulk submission of large numbers of annotated DNA sequences without the need for manual data editing. To demonstrate the utility of `annonex2embl`, I employ the software to automatically prepare a dataset of more than 1,500 fungal DNA sequences for submission to ENA.

## MATERIALS AND METHODS

### Overall design

`annonex2embl` was written in Python (Python Software Foundation 2019) and is compatible with both Python versions 2 and 3. The software reads aligned DNA sequences and a set of sequence annotations from an MSA, splits the MSA into individual sequences, adjusts the annotations to the length of the individual sequences, connects each sequence to the correct metadata co-supplied in a data spreadsheet, and converts each sequence plus its associated annotations and metadata into a sequence record of an EMBL-formatted multi-record output file. The software is exclusively controlled via command-line parameters to facilitate its integration into automated bioinformatic pipelines, but also maintains full functionality as a stand-alone tool. The internal integrity of the software is confirmed via a series of unit tests (Pajankar 2017) which evaluate the proper construction of sequence records, feature tables and gene annotations, the length equilibration of individual sequence records, the automated search of gene product names, and the parsing and transfer of metadata to the sequence records. A display of the internal design of `annonex2embl` is given in Supplementary Figure S1.

### Input and output

To execute `annonex2embl`, a minimum of six types of input information must be supplied by the user: (i) an MSA containing two or more aligned DNA sequences, (ii) annotations for the aligned sequences, (iii) a data table containing metadata for the sequences, (iv) a one-line description for the genomic region represented by the sequences, (v) the name of the output flatfile, and (vi) the name and email address of the person preparing the sequence submission. The MSA and the sequence annotations must be supplied as different blocks within a single NEXUS file, which can store sequence alignments, character sets, and phylogenetic trees as separate blocks of information (Maddison et al. 1997). Specifically, the MSA must be supplied as a ‘data’ block, the sequence annotations as an accompanying ‘sets’ block of the same NEXUS file. Multiple independent character sets can be specified per sets block, each of which defines the title, type, and position of an annotation feature across all sequences of the MSA (see the ‘sets’ block of ‘input 1 of 2’ in Figures 1 and 2). Every character set title must follow a specific format: it must encode - in that order - the name, the type and, optionally, the reading direction of the annotation feature, each separated by an underscore. The specification of the feature type is restricted to the vocabulary of feature keys defined by the INSDC (http://www.insdc.org/files/feature_table.html#7.2; accessed on 22-Sep-2019). The specification of the reading direction is limited to the keywords ‘forward’ and ‘reverse’, with the 5’ to 3’ direction (‘forward’) representing the default value. For example, the hypothetical coding sequence ‘foo’ with a forward reading direction would be represented by either character set definition `foo_CDS` or `foo_CDS_forward` (see the first character set of the ‘sets’ block of ‘input 1 of 2’ in Figures 1 and 2). Different character sets may overlap or even be identical in position with other character sets, and may be of any length ≥1. The metadata table must be supplied as a comma-delimited spreadsheet and contain a minimum of two data columns (‘input 2 of 2’ in Figure 1). The first column (titled ‘isolate’ by default) connects the individual rows of the metadata table to the DNA sequences of the MSA; its values must be identical to the sequence names of the MSA. The second column (titled ‘organism’ by default) contains the taxon names of the organisms represented by the sequences; its values must be identical to scientific taxon names registered at the NCBI taxonomy database (Federhen 2012). Next to these mandatory columns, the metadata table may comprise any number of additional data columns as long as the column titles are limited to the vocabulary of source feature qualifiers defined by the INSDC. The description of the genomic region represented by the MSA must be supplied as a short alphanumeric character string and characterize the entire genomic region delineated by the sequences (e.g., ‘gene1 complete sequence, and gene2, partial sequence’). `annonex2embl` adds this description as part of the official description line to each sequence record of the resulting flatfile. The name and email address of the person preparing the sequence submission must be supplied as separate character strings and are necessary for both, their inclusion in the resulting flatfile as well as to enable optional features of `annonex2embl` such as the online gene product lookup. Next to this mandatory input, several optional input parameters can be specified to control specific processes during software execution.

**Figure 1.**
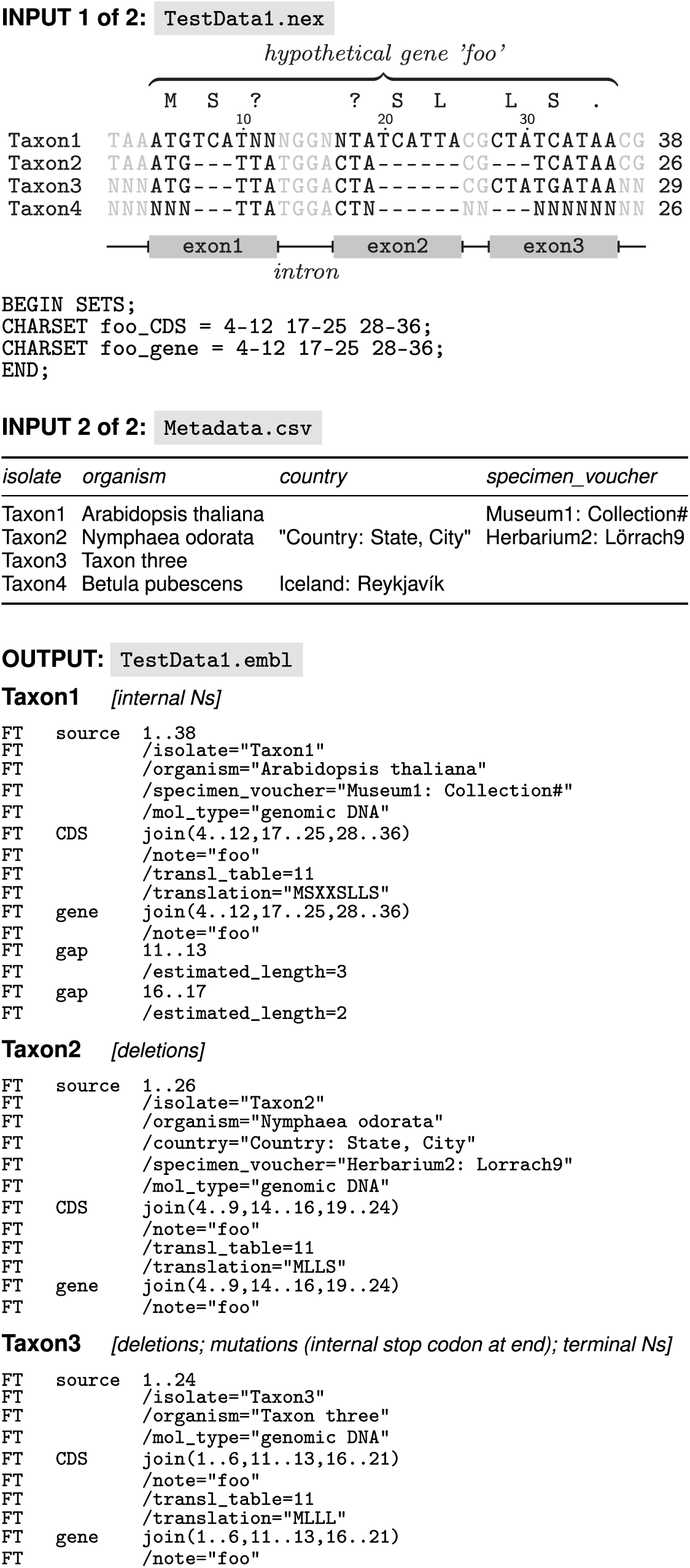
Schematic example of the input passed to, and the output received from, `annonex2embl` under test dataset 1. Only the matrix of the data block in the NEXUS input file is visualized. The country value for taxon 2 in the metadata input table is enclosed by double quotation marks because it contains a comma. Taxon 4 is not saved to the output by `annonex2embl` because the number of unambiguous nucleotides in its sequence is smaller than the minimum number required by the automated validation conducted by Webin. The sequence translation above the MSA refers to the sequence of taxon 1, is not part of the actual input, and provided to assist readers in relating the input to the output information.

**Figure 2.**
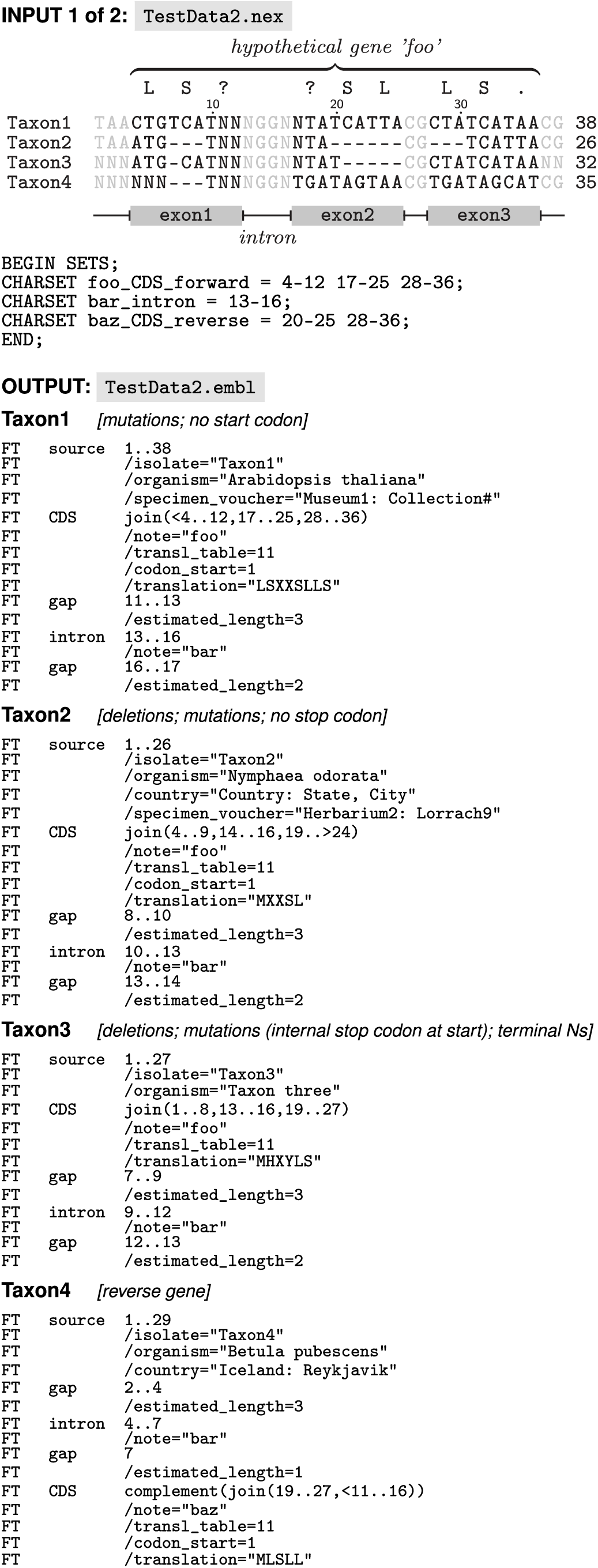
Schematic example of the input passed to, and the output received from, `annonex2embl` under test dataset 2. Input 2 of 2 is the same as in Figure 1 and, thus, not displayed. To conserve space, the source qualifier `/mol_type`=`“genomic DNA”` was removed from each output feature table.

Based on the aggregate of all input information, `annonex2embl` reads and parses the aligned DNA sequences and their annotations from the NEXUS file, as well as the sequence metadata from the comma-delimited spreadsheet, interlaces annotations, metadata, submitter info and general sequence description with the individual sequences, and saves of each DNA sequence plus its associated annotations and metadata one by one in the EMBL flatfile format. The output of `annonex2embl` is a multi-record EMBL flatfile as well as an accompanying manifest file. The flatfile comprises a set of individual sequence records whose number should be equal to the number of DNA sequences in the input MSA. If the number of sequence records is lower than the number of sequences in the MSA, not all of the aligned sequences were processed successfully. `annonex2embl` informs the user of any issues encountered but does not necessarily terminate the overall processing of the input MSA. This design allows the rapid and efficient conversion of a large number of aligned DNA sequences, even if some of them exhibit incorrect specifications and require a separate treatment. Moreover, `annonex2embl` automatically skips the processing of all input sequences with a length of ten or fewer unambiguous nucleotides (e.g., taxon 4 in Figure 1), as such sequences do not pass the automated validation procedure of the Webin submission portal. The manifest file specifies the name, location, and associated study number of the flatfile and is required for the command-line driven submission of the flatfile via Webin-CLI. A set of example input and output files that illustrate the conversion process conducted by `annonex2embl` is co-supplied with the package (folder ‘examples’) and portrayed as Figures 1 and 2.

All input files required for the execution of `annonex2embl` can be generated via standard biological software applications. The NEXUS file, for example, can be generated by various user-friendly sequence alignment editors (e.g., PhyDE, Müller et al. 2010) as well as via the application of biological script libraries (e.g., BioPython, Cock et al. 2009). The metadata table can be generated by any common spreadsheet editor. Users can compile a wide variety of information in the metadata table for all or only some of the DNA sequences of the MSA as long as the sequence IDs and the organism names are present for all entries. Placeholders for empty table cells are unnecessary, as `annonex2embl` automatically removes vacant cells upon sequence processing.

### Adherence to INSDC conventions

`annonex2embl` strictly adheres to the naming and formatting conventions defined by the INSDC for feature keys and their qualifiers. The INSDC has defined a list of 52 feature keys and 102 associated qualifiers to represent functional annotations in DDBJ, EMBL and GenBank sequence flatfiles (feature table definition version 10.8, Karsch-Mizrachi et al. 2018). Any EMBL-formatted flatfile intended for submission to ENA must adhere to this terminology and the associated formatting rules to pass the automated validation procedure of the Webin submission portal (Gibson et al. 2015). By extension, any software tool that aims to convert DNA sequences to EMBL flatfiles must also enforce these conventions. Without adherence to these rules, users would need to post-process the resulting flatfiles to ensure compatibility with the EMBL flatfile standard, which can be prohibitively time-expensive. `annonex2embl` enforces the INSDC conventions by checking if all column names of the metadata table abide by the controlled vocabulary and, thus, match permissible qualifier names. For example, metadata information on the collection date and the collection locality of a sequenced organism must be named after, and formatted according to, the INSDC source feature qualifiers ‘collection_date’ and ‘country’, respectively. Users are, thus, compelled to employ the INSDC-approved terminology and its formatting rules when compiling the input data. Specifically, the character set names of the sequence annotations, as well as the column names of the metadata table, must conform to the approved terminology, and the content of the metadata table must adhere to the formatting rules. Similarly, `annonex2embl` enforces the correct character encoding for all metadata content. The INSDC has specified that feature qualifier values may only be composed of printable ASCII characters. Consequently, `annonex2embl` automatically replaces all non-ASCII characters in the metadata with ASCII-compatible characters before integrating the character strings as qualifier values in the feature table (see, for example, specimen voucher information of taxon 2 of ‘input 2 of 2’ in Figure 1 and compare it with the corresponding feature table line in the output section of the same figure). Likewise, `annonex2embl` transforms all user-supplied sequence annotations to appropriate feature table entries by converting the character set definitions to INSDC-approved feature keys with fitting qualifiers. Taken together, the strict adherence by `annonex2embl` to the naming and format conventions of the INSDC streamlines the generation of EMBL flatfiles without the need for manual post-processing.

### Installation and usage

`annonex2embl` is accessible via the Python package index under http://pypi.python.org/pypi/annonex2embl and can be installed using the command `pip install annonex2embl`. It has been successfully tested on Linux (Arch Linux 4.18, Debian 9.9, and Ubuntu 16.04), MacOS (macOS 10.13), and Windows (Windows Server version 1803) under both Python 2.7 and 3.7.

The usage of `annonex2embl` is exclusively controlled via command-line parameters. At a minimum, a user must enter six input parameters: the name of, and file path to, a NEXUS file (command-line parameter `-n`); the name of, and file path to, a comma-delimited metadata table (`-c`); a quotation-mark enclosed text string that describes the genomic region represented by the DNA sequences (`-d`); the name of, and file path to, the output flatfile (`-o`); and the name (`-a`) and email address (`-e`) of the person preparing the sequence submission, both as quotation-mark enclosed text strings. In addition to these mandatory parameters, users may invoke up to eleven optional parameters. For example, users may wish to have `annonex2embl` test if each taxon name supplied via the metadata has been registered as a scientific taxon name with the NCBI taxonomy database (command-line parameter `--taxonomycheck`). ENA does not accept the submission of DNA sequences representing taxa that have not been registered with the NCBI taxonomy database, rendering an *a priori* evaluation of their registration critical. Similarly, users may wish to have the software automatically search and add standardized gene product names to the feature tables of the output flatfile (`--productlookup`), which is important for a complete representation of the DNA sequences (Pirovano et al. 2017). When selecting this option, `annonex2embl` parses the gene abbreviations from the character set definition titles of the sequence annotations, communicates with NCBI to identify the complete gene names, and adds appropriate product qualifiers to the relevant feature keys. For example, the gene abbreviation ‘matK’ would be identified as, and linked to, the complete gene name ‘maturase K’, which, in turn, would be added to the qualifier ‘product’ of the feature key ‘gene’. The full set of available input parameters, their default values, and a short explanation of each parameter can be displayed by invoking the help command via `annonex2embl -h`.

To invoke `annonex2embl` on one of the example input files co-supplied with the package, a user may type the following command in a terminal:

~~~
annonex2embl \
-n examples/input/TestData1.nex \
-c examples/input/Metadata.csv \
-o examples/output/TestData1.embl \
-d “description of alignment here” \
-a “your name here” \
-e “your_email_here@yourmailserver.com”
~~~

To communicate any issues encountered while processing the input data, `annonex2embl` prints short warning or error messages to the screen. Warning messages indicate the skipping of individual sequence records due to issues restricted to these records, whereas error messages indicate the termination of the software prior to processing the last record of a file. For example, `annonex2embl` would provide a warning message if an annotation feature of type ‘CDS’ was not added to the feature table of a sequence record if the underlying amino acid sequence contained a stop codon immediately after the indicated start codon (e.g., hypothetical coding sequence ‘baz’ in taxa 1, 2, and 3 in Figure 2). Similarly, the software would provide a warning message if a sequence record was not saved to the output flatfile if an organism name was not encountered in the NCBI taxonomy database (e.g., taxon 3 in Figures 1 and 2), assuming that the optional parameter for taxonomy checks had been selected.

Upon execution, `annonex2embl` generates a submission-ready EMBL-formatted flatfile as well as an accompanying manifest file, which is specifically required for command-line driven sequence submissions to ENA. `annonex2embl` does not, however, conduct the sequence submission itself. Instead, the submission of the flatfile to ENA remains the responsibility of the user. This design decision was taken for two reasons: First, with Webin-CLI, ENA offers an official command-line software tool for the upload and submission of annotated DNA sequences via the Webin submission portal, whose usage shall be encouraged. Second, users should always validate the accuracy and content of the flatfiles before a submission via the proper validation tools (Harrison et al. 2019), such as Webin-CLI or the stand-alone ENA flatfile validator (Gibson et al. 2015). Upon submission of a flatfile to ENA, the user receives a list of unique accession numbers from ENA that permanently identify the sequences.

### Application on empirical data

To demonstrate its utility, `annonex2embl` is employed to prepare a large-scale dataset of annotated fungal DNA sequences for submission to ENA. Specifically, the software is used to automatically convert an annotated MSA in NEXUS format comprising a total of 1,518 individual DNA sequences, and a corresponding metadata table, into a submission-ready EMBL-formatted flatfile. The DNA sequences were collected in an investigation on fungal molecular diversity of the Canarian archipelago (Gruenstaeudl et al. 2013) and have not been submitted to any sequence database. Upon conversion to an EMBL flatfile, the sequences are first validated and then uploaded to ENA, both via Webin-CLI v.1.8.11. The run-times of both `annonex2embl` and Webin-CLI are measured and compared. The input files for this test submission are available as references for format and content from Zenodo at https://zenodo.org/record/3517124.

## RESULTS

Using `annonex2embl` to prepare a total of 1,518 aligned DNA sequences for submission to ENA was found to be rapid and reliable. Specifically, the software required 1.4 minutes for converting the annotated MSA and the co-supplied metadata into a submission-ready EMBL-formatted flatfile when only the mandatory parameters for a submission via the Webin portal were employed (Table 1). Thus, the conversion was an order of magnitude faster than the subsequent flatfile validation or the subsequent flatfile upload to ENA, both of which were conducted via Webin-CLI and required 15.9 minutes and 19.4 minutes, respectively. No data processing other than the invocation of `annonex2embl` was required before validating and uploading the flatfile, and not a single sequence record required post-processing, illustrating that `annonex2embl` can prepare a large number of annotated DNA sequences for submission to ENA in an automatic and streamlined fashion. The submitted fungal DNA sequences are available on ENA under accession numbers LR730418-LR731935.

**Table 1.**
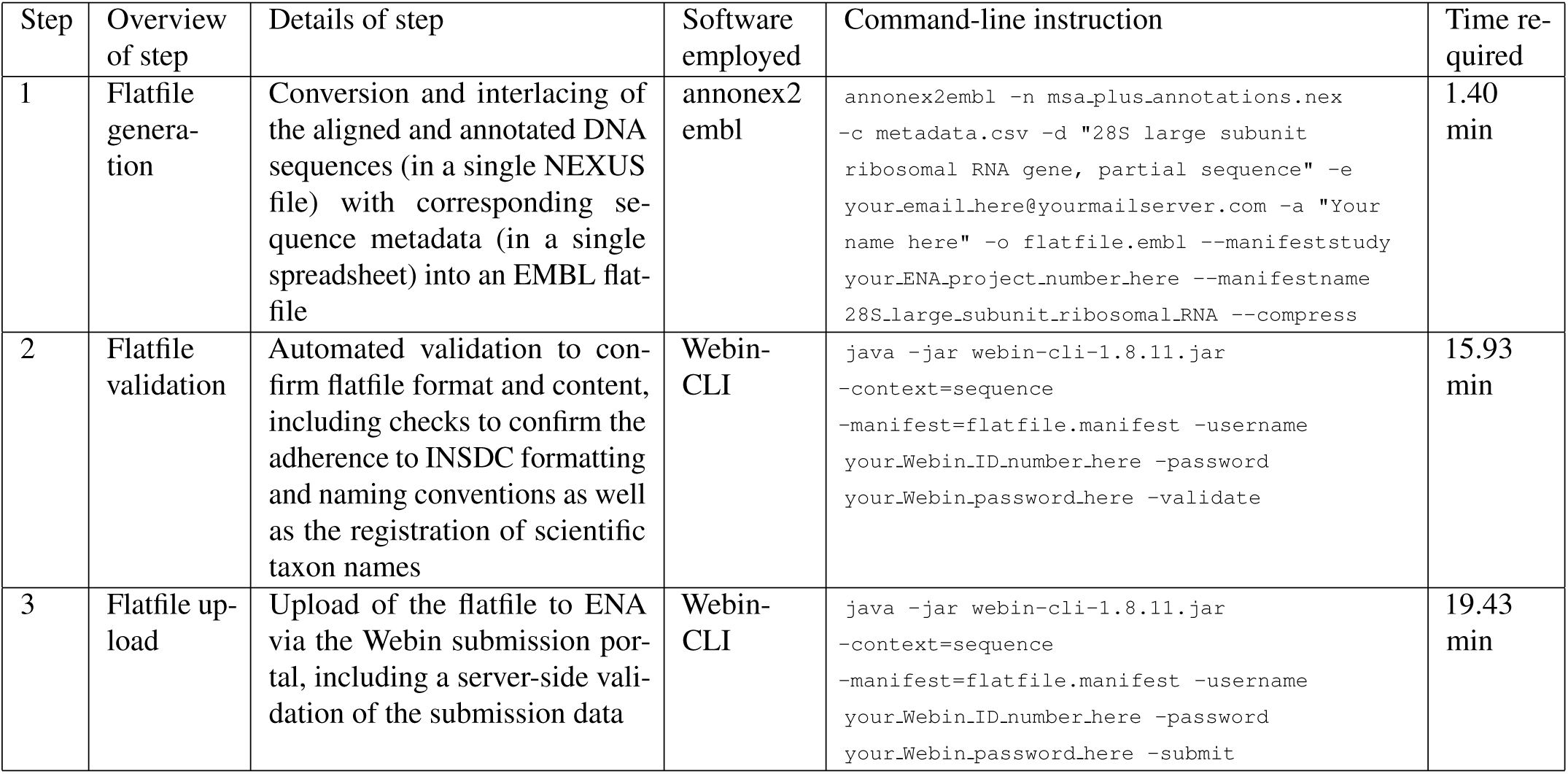
Overview of, the time required for, and the command-line instructions used in, the bioinformatic steps involved in the bulk preparation, validation, and submission of 1,518 fungal DNA sequences to ENA via `annonex2embl` and the command-line route of the Webin submission portal. Command-line instructions refer to usage on the Bash shell under Linux.

Moreover, `annonex2embl` has been employed by several molecular phylogenetic investigations to facilitate their sequence submissions to ENA. Specifically, the software has so far been used in Roy et al. (2017), Korotkova et al. (2018), Canal et al. (2018), Borsch et al. (2018), and Falcon-Hidalgo et al. (2019) to submit DNA sequence data via the Webin submission portal.

## DISCUSSION

Several routes for preparing and submitting annotated DNA sequences to ENA exist. First, researchers can prepare annotated DNA sequences for submission as pre-formatted data spreadsheets (so-called ‘checklists’), which are manually filled by the user with nucleotide sequences and associated metadata and then uploaded to ENA via the interactive route of the Webin submission portal (Silvester et al. 2018). Idiosyncrasies of the genomic regions necessitate the application of different checklist types, rendering the submission of diverse sets of sequences time- and labor-expensive. However, a recently developed software tool enables the automatic conversion of DNA sequences to some of the more commonly applied checklist types (Gruenstaeudl and Hartmaring 2019) and may, thus, be an option for small-scale submissions. Second, researchers can prepare annotated DNA sequences for submission as EMBL-formatted flatfiles, which can be uploaded to ENA via the command-line route (Harrison et al. 2019) or the programmatic route (Silvester et al. 2018) of the Webin submission portal. Programmatic submissions require that input flatfiles are accompanied by XML authentication and submission files which are difficult to generate for nonexperts. Hence, the command-line driven route of constructing, validating, and uploading EMBL flatfiles currently represents the most efficient and user-friendly method for large-scale sequence submissions to ENA.

Only few software applications exist that can generate EMBL-formatted flatfiles. Most of these tools operate only on individual sequences or require additional file conversions for their output to conform to the EMBL flatfile standard, rendering them unsuitable for automated, large-scale submission preparations. For example, the open-source genome viewer Artemis (Rutherford et al. 2000) can save annotated DNA sequences as flatfiles that abide by the EMBL flatfile format, but can only operate on one sequence at a time. Moreover, functional annotations must be added for each sequence individually in this viewer, rendering sequence submissions prohibitively time-expensive for all but the smallest datasets. Similarly, the commercial software suite Geneious (Kearse et al. 2012) can save DNA sequences in the GenBank flatfile format, which must be further converted to the EMBL flatfile format via additional software tools (e.g., EMBOSS Seqret, Olson 2002). While Geneious allows the semi-automatic propagation of annotations across aligned sequences, it is primarily driven via a graphical user interface, effectively precluding its integration into automated bioinformatic pipelines. Moreover, the conversion route enabled by Geneious is dependent on the continued compatibility of multiple, independent software tools and requires users to afford the licensing cost for the software suite. Similar functionality as provided by Geneious in conjunction with file converters can be achieved via the legacy NCBI tool Sequin (Benson et al. 2013), which can also transfer annotations across DNA sequences but, in like manner, requires additional conversion steps to generate EMBL-formatted flatfiles. Next to these stand-alone software tools, several online services offer the automatic submission of DNA sequences to ENA; however, the functionality of these web services is difficult to assess, as many of them are inaccessible in their original form (e.g., CDinFusion, Hankeln et al. 2011, attempt to access on 24-Oct-2019) or have not made their source code and, thus, the details of their underlying data processing publicly available. `annonex2embl`, by contrast, represents an open-source software tool that is transparent in its underlying processes, easily customizable, and available to users without charge. It enables the automated, command-line driven processing of large-scale sequence datasets directly into EMBL-formatted flatfiles and can be easily integrated into automated bioinformatic pipelines.

With the application of `annonex2embl`, users can prepare bulk submissions of DNA sequences without the need for additional data processing, particularly regarding the assignment of sequence annotations. Previously, researchers had to invest considerable time to manually prepare their sequence data for submission to public sequence databases (Pirovano et al. 2017; Gruenstaeudl and Hartmaring 2019). By employing `annonex2embl`, users can harness the advantages inherent to aligned sequences for their automated, streamlined conversion to records of a submission-ready flatfile because aligned sequences are uniform in length, making them amenable to automatic processing. For example, by defining annotations on aligned sequences, the annotations do not need to be specified one sequence at a time, but can be simultaneously assigned across all input sequences. Specifically, the annotations can be propagated to each sequence of an MSA due to the positional homology among them. Such a bulk assignment provides greater consistency between annotations and is an efficient means for preparing large-scale submissions. However, the bulk assignment of annotations on aligned sequences is only possible as long as the implicit length differences among the individual sequences are accommodated. MSAs typically comprise sequences of various length, with spaces inserted within sequences to enable the alignment of homologous nucleotides (Morrison et al. 2015). Accounting for these length differences while maintaining correct annotation assignments is a complex process and becomes especially intricate when the differences are the result of overlapping but non-identical insertions and deletions. `annonex2embl` maintains correct annotation assignments while accounting for implicit length differences among the sequences by employing a process that automatically transfers all length changes of the sequences to the location positions of their annotation features, thus keeping them in sync.

Another advantage of using MSAs for preparing bulk submissions of DNA sequences to public databases is the streamlined addition of metadata to the individual sequences. Sequence records in public databases should contain as much metadata information as possible, allowing the cross-linking of the submitted data with, and its re-usability by, other analyses and, thus, enhancing its value to other researchers (Meyer et al. 2019). In particular, a detailed description of the identity and location of the organism from which a sequence was generated, references to biological collections that preserve its voucher specimens, and bibliographic information that link the sequence to the published description of the dataset are essential for the scientific re-use of these sequences (Kans and Ouellette 2001). Preparing large-scale sequence sets for submission to public sequence databases should, consequently, include an automated interlacing of metadata to each sequence record, which `annonex2embl` implements via a sequence-specific data integration from a co-supplied metadata table.

In summary, `annonex2embl` enables the automated preparation of annotated DNA sequences plus associated metadata for bulk submission to the online sequence archive ENA and will likely accelerate the flow of novel sequence data to this archive, particularly for sequences from phylogenetic and population genetic investigations.

## Acknowledgments

The author thanks Gabriele Dröge of the Botanischer Garten und Botanisches Museum Berlin for valuable discussions as well as Yannick Hartmaring and Nils Jenke of the Freie Universität Berlin for assistance with testing and debugging the software.

