## Supplementary Data for "annonex2embl: automatic preparation of annotated DNA sequences for bulk submissions to ENA"

Michael Gruenstaeudl[1*]

[1] Institut für Biologie, Freie Universität Berlin, 14195 Berlin, Germany  *

**Supplementary Fig. S1** Display of the internal design of annonex2embl. Double-circled fields indicate input and output of the software, dashed arrows and fields indicate optional parameters and processes. All optional input parameters are listed in the right-most column for easier viewing but are provided upon the initiation of the software (i.e., top row).

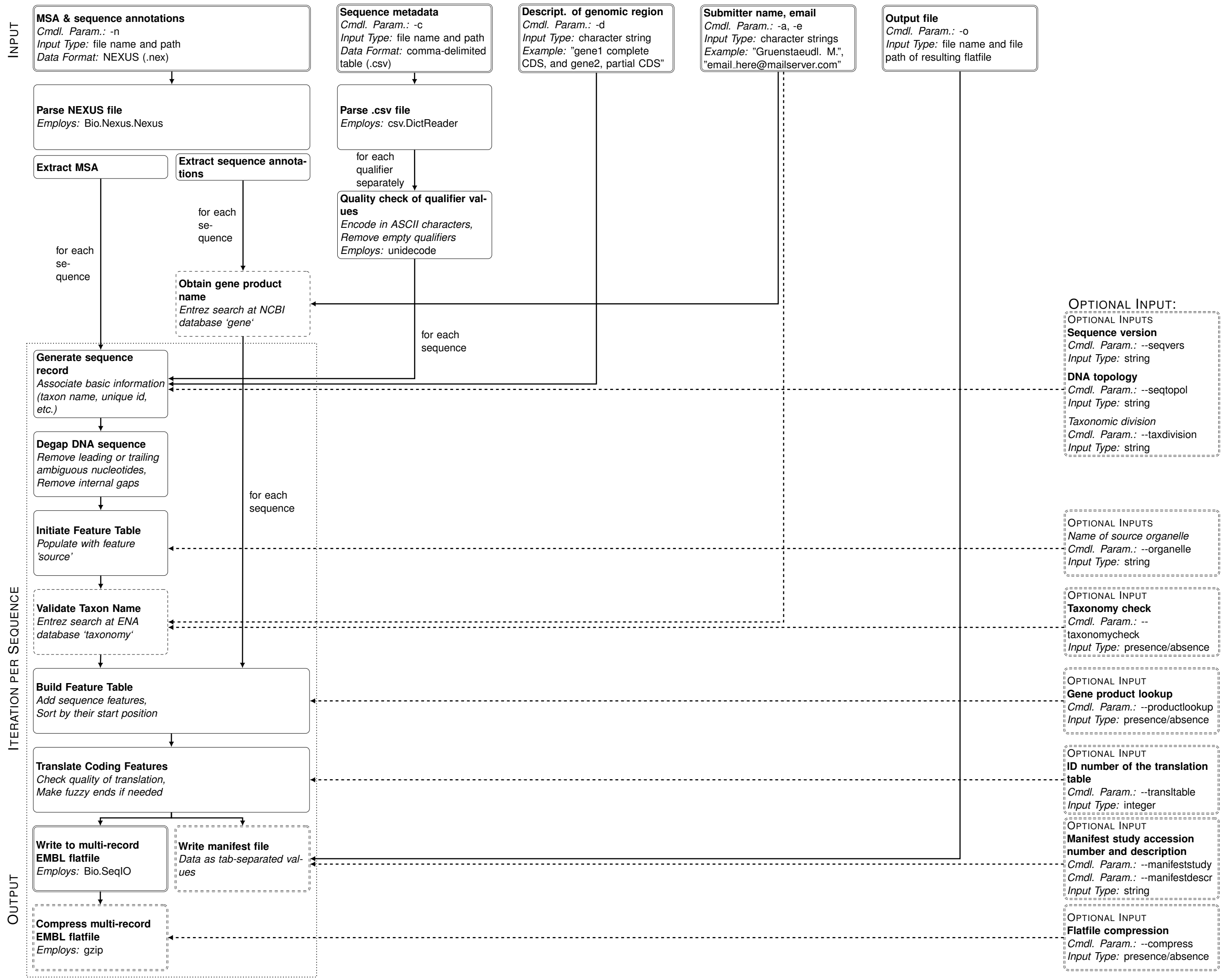
